# Firing-rate-modulated spike detection and neural decoding co-design

**DOI:** 10.1101/2023.01.10.523472

**Authors:** Zheng Zhang, Timothy G. Constandinou

## Abstract

**Objective:** Translational efforts on spike-signal-based implantable brain-machine interfaces (BMIs) are increasingly aiming to minimise bandwidth while maintaining decoding performance. Developing these BMIs requires advances in neuroscience and electronic technology, as well as using low-complexity spike detection algorithms and high-performance machine learning models. While some state-of-the-art BMI systems jointly design spike detection algorithms and machine learning models, it remains unclear how the detection performance affects decoding.

**Approach:** We propose the co-design of the neural decoder with an ultra-low complexity spike detection algorithm. The detection algorithm is designed to attain a target firing rate, which the decoder uses to modulate the input features preserving statistical invariance.

**Main results:** We demonstrate a multiplication-free fixed-point spike detection algorithm with nearly perfect detection accuracy and the lowest complexity among studies we have seen. By co-designing the system to incorporate statistically invariant features, we observe significantly improved long-term stability, with decoding accuracy degrading by less than 10% after 80 days of operation. Our analysis also reveals a nonlinear relationship between spike detection and decoding performance. Increasing the detection sensitivity improves decoding accuracy and long-term stability, which means the activity of more neurons is beneficial despite the detection of more noise. Reducing the spike detection sensitivity still provides acceptable decoding accuracy whilst reducing the bandwidth by at least 30%.

**Significance:** Our findings regarding the relationship between spike detection and decoding performance can provide guidance on setting the threshold for spike detection rather than relying on training or trial-and-error. The trade-off between data bandwidth and decoding performance can be effectively managed using appropriate spike detection settings. We demonstrate improved decoding performance by maintaining statistical invariance of input features. We believe this approach can motivate further research focused on improving decoding performance through the manipulation of data itself (based on a hypothesis) rather than using more complex decoding models.

## 1. Introduction

Brain-machine interfaces (BMIs) have shown great potential for providing new treatment for disorders [1, 2], rehabilitation [3, 4] and human-computer interaction [5, 6]. Such amazing achievements are the result of the joint effort of neuroscience, the development of neural signal processing and artificial intelligence, and advanced recording technology. Designing minimally invasive wireless implantable BMIs with high channel counts is the current trend in next generation BMIs [7, 8]. High channel count implants can provide high-resolution recordings that provide the approach for improving the decoding performance and reliability. Through miniaturisation and wireless connectivity, there is no need for percutaneous wires, ensuring that the BMIs are the most portable with the least risk of brain damage or infection.

Next generation BMIs pose significant new challenges. On the implant end, enormous data is collected with increased channel counts. Bandwidth reduction and lightweight on-edge processing algorithms are essential to guarantee that the on-implant processing and transmission power is within the power budget. This prolongs the battery’s lifetime whilst preventing prevents damage to brain tissue. On the back end, the decoding performance can be affected by implant degradation. Improving the long-term stability of neural decoding is another essential requirement for invasive BMIs. Designing advanced decoding models [9, 10] and utilising long-term stable features are two keys to preventing performance degradation [10–12]. Multi-Unit Activity (MUA) is one feature that can fulfil mentioned requirements. It has better long-term stability and consumes less computation to be extracted compared to the Single-Unit Activity (SUA), which is widely used in practice [6, 13, 14].

For MUA-based BMIs, on-implant spike detection is often combined with off-implant decoding. Such systems however typically only utilise the most basic spike detection methods (e.g. fixed threshold defined by training statistics).

The system co-design must take into account different requirements/constraints for the spike detection algorithm and the neural decoder. The spike detection algorithm needs to be designed to minimise hardware complexity and power consumption as this will be performed on the implant. In contrast, the neural decoding will operate on an external node with more resources and a larger battery, such as a mobile phone or wearable device, where the computational requirements are more relaxed in order to improve decoding accuracy.

The co-design consists of an ultra-lightweight adaptive spike detection and a neural decoder with improved long-term stability addressing requirements mentioned previously. Using the proposed system co-design, we also investigate the relationship between spike detection and neural decoding performance in real recordings where no ground truth is available. Three findings have been observed in this work.

1. Detecting small spike peaks around the noise floor can improve the decoding performance. This reveals that there is the neural activity far away that is informative relatively. Utilising this extra activity in decoding can improve decoding performance other than being detrimental despite detecting additional noise that is often considered to be detrimental. Therefore, applications that target maximising decoding performance in the realm of neuroscience should follow this approach where the threshold should be set only a little above the noise floor, i.e. sensitivity is more important than specificity.
2. Detecting only significant peaks causes less than 1% performance degradation. As fewer spikes are detected, less information needs to be transmitted. This can be the best trade-off between the data bandwidth and decoding performance, which is often preferable for wireless brain machine interfaces.
3. Long-term stability is degraded through inconsistent feature representation. Normalising the input feature with the training set statistics is a standard step before feeding data into neural networks. However, training set statistics are no longer valid as recording quality degrades. We found that normalisation using such statistics causes inconsistent feature representation, which is one key reason for long-term decoding degradation. Performance degradation can be reduced by keeping track of the feature statistics instead of using training data statistics, and distinguishable improvements can be made using the proposed spike detection and decoding co-design system. Using a broader spatial distribution of neural activities, i.e. spike signals further away, also benefits long-term stability.

This paper is structured as follows: Section 2 details the challenges in both spike detection and neural decoding. Section 3 introduces the datasets used in this paper. Section 4 introduces the proposed co-design system, including the novel firing-rate-based spike detection algorithm and input-consistent decoding model. Section 5 shows the performance of the proposed spike detection algorithm on the synthetic dataset, the corresponding hardware cost and The comparison to other algorithms. It also includes the relationship between spike detection and neural decoding, the decoding performance of the co-design system, and its improvement in long-term stability. Section 6 discusses the concern on data consistency in neural decoding and the opportunity this work provided to brain machine interface. Section 7 concludes this work.

## 2. Challenges in wireless invasive brain machine interfaces

Challenges in designing wireless invasive brain machine interfaces are still significant. One key challenge is power consumption. On the one hand, a massive amount of data is collected on the implants. The recording instrumentation together with processing and transmission can consume enormous power. On the other hand, the brain tissue around the implants is hugely sensitive to temperature increases and vulnerable, which can be damaged by only a 1 ^*◦*^*C* temperature increase [15]. Advanced low-power hardware design is one key factor in overcoming this challenge while reducing the data bandwidth also plays an important role.

### 2.1. Neural spike detection

Feature extraction is essential for reducing the data bandwidth and keeping the implant power within budget. Using the MUA signal, the bandwidth can be reduced from more than 300 kbps/channel to 1kbps/channel/neuron firing around the implanted probes [16, 17], and even as low as 30bps/channel after compression [18]. In the meantime, useful information can be mostly preserved as spikes are the most informative within intracortical signals.

In order to obtain MUA signals, spike detection is commonly used. Spike detection applies a threshold to the recordings, and for the duration of the signal is above the threshold, spikes are recognised. Various spike detection algorithms have been proposed using the wavelet transform [19–21], control theory [22, 23], robust estimation of noise and spike distributions [24–26], and machine learning [27]. These algorithms can detect spikes with excellent accuracy. However, their computation requirement can be excessive for on-implant use, even though some have already been optimised towards low-power implementation. A more simple form of spike detection using nonlinear energy operator (NEO) [28, 29], amplitude slope operator [30, 31] to emphasise the spikes, and defining the threshold based on gross signal statistics can be ideal for on-implant use due to their simplicity.

Another aspect that becomes increasingly important is real-time processing and algorithm adaptiveness. Real-time signal processing is essential for BMI applications guaranteeing the fast response of the whole system, and adaptive spike detection enables the threshold to adapt to the varying brain environment in time and across channels or electrodes. Various techniques have been used, for example. In [30], bit shifts are used to replace multiplication, improving the real-time system response time. In [32], dual thresholds derived from smoothed and non-smoothed NEO is used to guarantee high accuracy in both low and high SNR level to improve adaptiveness.

### 2.2. Challenges in real-time on-implant spike detection

Several challenges exist, and the trade-off between algorithm complexity and performance is still unresolved. Low-complexity spike detection algorithms can consume little power, which is preferred for wireless implantable BMIs. In [33] and [24], different low-complexity spike detection algorithms have been tested on FPGA or ASIC, addressing such a trade-off. However, the adaptiveness of the evaluated statistical-based approach is of concern. In [34], it is reported that utilising a threshold based on noise statistics can never establish a threshold with consistent performance across different noise levels, while jointly using noise and spike statistics does. However, spike peak statistics are difficult to be estimated as spikes are unknown before they can be detected. In addition, extra computation is needed.

We proposed a firing-rate-based algorithm in [35] based on the fact that neurons in a certain brain region are firing at a normal rate. If the average detection rate is close to that normal rate, the current threshold can be considered to be a reasonable threshold. We developed several techniques enabling the target spike rate to update to the correct rate adaptively, eliminating the effect of inaccurate initial settings. The proposed online method achieves better detection performance than the offline thresholding algorithms. At the same time, the hardware cost is even less than the threshold algorithms based on calculating the local average.

Another challenge is in evaluating the spike detection algorithms. There is generally no spike ground truth in neural recordings unless with manual labelling. Though some studies tried to design an auto-labelling algorithm [36], new metrics [37] or use intro-extracellular-paired recordings [38], most of the spike detection algorithms are evaluated using synthetic data [24, 30, 39–44]. There are considerable differences between the synthetic data and real recordings w.r.t noise level, signal variety, etc. It is not guaranteed that the performance of a spike detection algorithm can be maintained in practice.

In the context of the MUA-based brain machine interface, the output of spike detection is the input of the neural decoding. An even more significant and under-investigated challenge is to design a spike detection algorithm that improves the neural decoding performance rather than solely focusing on the spike detection performance. These two elements are always optimised individually in isolation. To the best of our knowledge, no work exists in the literature that optimises spike detection and neural decoding together holistically considering overall system performance.

### 2.3. Spike-based neural decoding and its challenges

MUA-based neural decoding has been widely applied in decoding subject movement[45– 47], speech[6, 48, 49], and vision[50, 51]. Previously, Kalman filters were the most widely used decoding model[52–54]. With the development of deep learning, Recurrent Neural Networks (RNN), Long-short Term Memory (LSTM), and the variations become more and more popular[6, 45, 46, 48, 49]. More recently, Generative Adversarial Networks (GAN) is used to overcome the difficulty of lacking train data[55], and attention mechanisms start to be used in neural decoding[56].

Though great progress has been made in neural decoding, most of the input features (spikes) are extracted using a relatively simple thresholding method via manually selected multipliers to signal statistics. Such an approach could be practical when enough memory is available to allow for statistically significant signal dynamics to be estimated. However, this can be unreliable in low-power design with constrained resources available on implants. It is commonly accepted that data quality is the essential factor in the success of machine learning or deep learning. This is especially so since the detection is effectively deciding what data is discarded. Therefore, it is again crucial to understand the relationship between spike detection and neural decoding performance and apply more advanced spike detection techniques to enhance the feature quality.

Long-time stability is also a challenge in neural decoding, as mentioned before. With the electrode ageing, scar tissue growth and foreign body response, the signal-to-noise ratio can drop as time goes by. Such signal degradation lowers the feature quality of the decoding models and limits the model’s capability in the long term. There are major efforts investigating the underlying mechanisms aiming to improve long-time stability [9–12]. Among different types of neural features, single-unit activities and spike waveforms are least robust in the long term, while multiunit activities and local field potentials are better [12, 45]. However, there is still no well-understood reason why different features can have varying effects on long-term stability and the commonly-accepted way to reduce the degradation.

## 3. Dataset

This study used two datasets, one synthetic and one realistic recording dataset to evaluate spike detection and neural decoding performance.

The synthetic dataset was first published in [39] and has subsequently been widely used in [24, 30, 40–44] to evaluate the spike detection performance. This dataset consists of four groups of recordings, namely Easy1, Easy2, Difficult1 and Difficult2. Each group consists of four different noise levels from 0.02 to 0.5 with a step of 0.05 in total 16 different pieces. The noise level in this dataset is defined with the ratio between the noise standard deviation and the average spike amplitude. The recordings are sampled at 24 kHz, and each consists of three types of spikes firing at 20 Hz (60 Hz in total) following Poisson distribution. More details of this dataset can be found in [39]. In our experiment, we used 60 s of each recording downsampled at 7kHz in fixed point representation as [30, 57] suggested

The data with real recordings was collected by Sabas Lab [58]. Utah arrays were placed into the primary motor cortex of a nonhuman primate. The data are sampled at 24414Hz, downsampled three times and rounded to fixed point representation. Its two-dimension hand movement was recorded with the brain signal while the free-moving subject spontaneously used a ticker to control the cursor reaching desired locations on the screen. The spike timing is provided in this dataset using a threshold derived from 3 to 5 times the signal rooted-mean-square (RMS) values. This dataset has been widely used for evaluating brain signal decoding algorithms [45, 59–61]. We have used the 24 pieces recording on Monkey Indy starting from 27^*th*^, June 2016 to 27^*th*^, January 2017, in total 5 hours.

In this work, instead of applying the decoding algorithm directly on the provided threshold crossings, we used the proposed spike detection algorithm to re-detect the spikes from the raw recordings. We evaluated our spike detection algorithm with decoding performance which can be translated to how much useful information is preserved after detection compared to the conventional spike detection algorithm. Meanwhile, we can observe the relationship between spike detection and decoding performance.

## 4. Spike detection and decoding system co-design

As shown in Fig.1a, the proposed system consists of signal acquisition, spike detection, spike binning and decoding, four modules. A target detection rate is set on both the spike detection and decoding model, modulating the input feature of the decoding model across different system settings and brain environments to maintain a consistent representation statistically.

**Figure 1.**
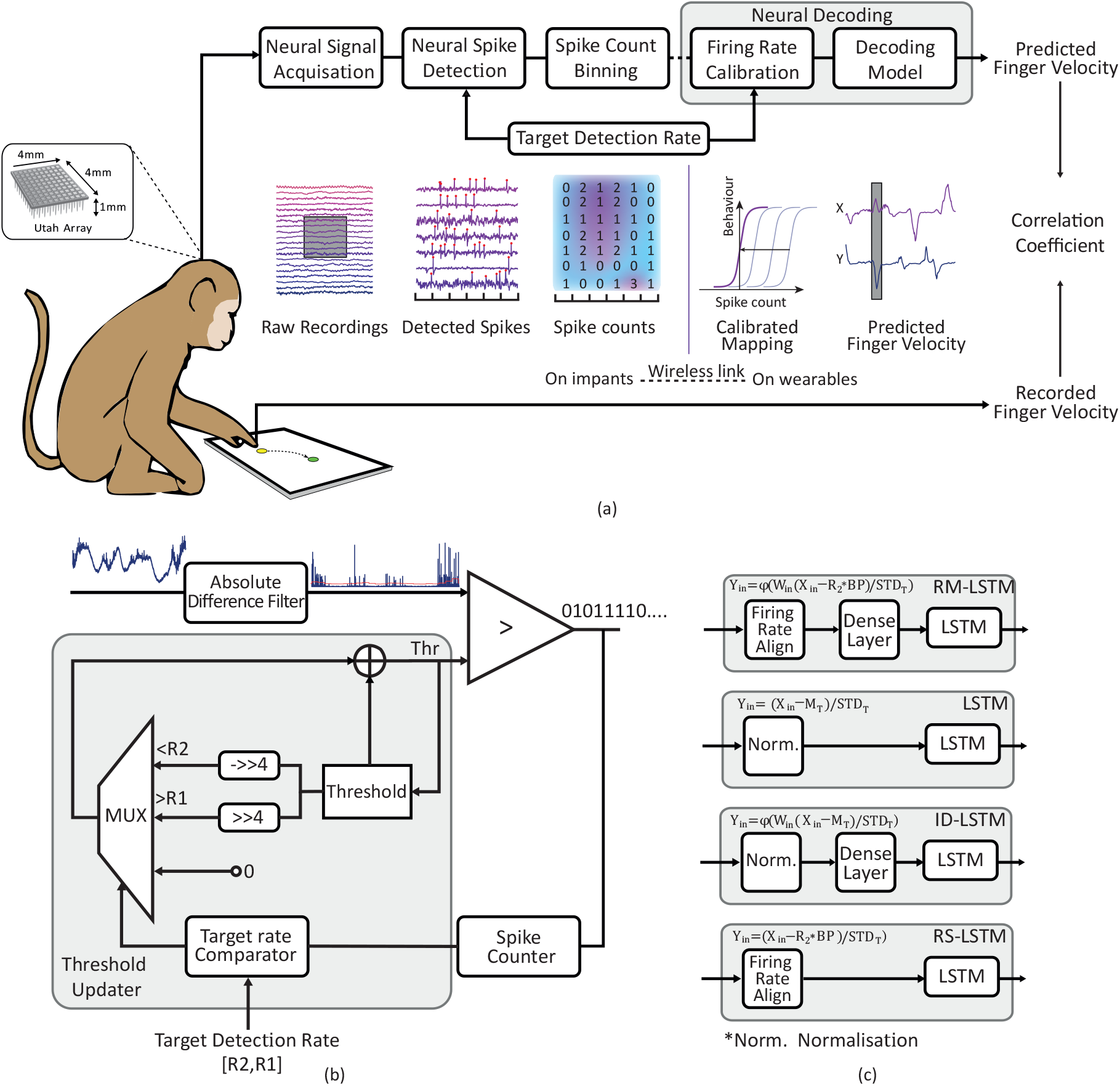
a) Spike detection - neural decoding system co-design. After digitising the intracortical signal, spikes are detected at the pre-set target detection rate. Detected spike counts are binned at a fixed period. Binned spike counts are then modulated using the target detection rate to align the feature representation over time. Finally, the decoding model predicts the target velocity, and its correlation coefficient to the real velocity is used to evaluate the system performance. b) Logic circuit of spike detection. The Absolute Difference Filter takes the input signal’s derivative and absolute value to remove the LFPs. The spikes are detected if the filtered signal outnumbers the threshold. The threshold is updated duty-cycled according to the relationship between the target detection rate and the number of detections counted by Spike Counter. Repeated detections are counted only once. c) Flowcharts of the neural decoding modules. All modules use the LSTM decoder but have different input layers. Rate-Modulated LSTM (RM-LSTM) subtracts the target detection rate from the binned spike counts followed by a dense layer. Compared to the proposed architecture, the conventional LSTM subtracts and divides the mean and standard deviation values calculated from the training dataset to normalise the input feature. Input Dense LSTM (ID-LSTM) consists of a dense layer with the standard normalisation. Rate Subtracted LSTM (RS-LSTM) only modulates the input feature using the target rate without the dense layer.

### 4.1. Firing-rate-based spike detection algorithm

#### 4.1.1. Preprocessing

The digitised raw intracortical neural signal can be offset by the local field potentials (LFP) and noise. In order to remove the LFP, we used the absolute value of the derivative following [35] as Equ.1 shows.

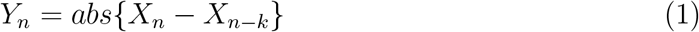

where *X*_*n*_ is the *n*^*th*^ input and *Y*_*n*_ is the corresponding output. *K* is sample rating dependent, and it is 2 for both datasets. Using such a simple absolute difference filter is effective enough in both datasets and significantly improves the signal SNR with minor computational cost.

#### 4.1.2. Firing-rate-based adaptive threshold

The proposed adaptive threshold is set based on the fact that the firing rate of a certain brain region is on average stable. In [62], researchers monitored twelve human motor neurons when the subjects were operating tasks with minimum effort, observing that the firing rate of each neuron is about 10Hz, and it became over 20Hz per neuron for maximum effort tasks.

Though the firing rate can fluctuate during the active or silent period, it is relatively stable in the long term. Therefore, by setting a reasonable target detection rate and threshold update strategy, we can control the threshold to saturate at the level, automatically detecting spikes at the desired rate.

Fig.1b shows the proposed spike detection algorithm. The target detection count interval [*R*_2_, *R*_1_], initial threshold *T*_0_ and updating period *P* is set at start. In one updating period, if *Y*_*n*_ *> T*_*n*_, a spike is detected, and 5 samples (in the 7kHz sample frequency case) are skipped to avoid multiple detections of one spike. The detection count *C* is increased by one. If *C > R*_1_, the threshold will increase by *p*%, and a new updating period is started. At the end of one update period, if *C < R*_2_, the threshold will decrease by *p*%. Otherwise, the threshold will be unchanged. In short, we set an acceptable detection number range [*R*_2_, *R*_1_], and the threshold will be decreased or increased if the detection rate is too low or high. Even though the initial threshold can be non-ideal, the threshold will eventually saturate at a level that fulfils the target. Here we set the *R*_1_ = 2*R*_2_. The value of *R*_1_ can vary and will be specified later in the result section.

*p*% determines the saturated speed and threshold resolution. A larger *p* means faster saturation but a less accurate threshold. Adaptively changing *p* can be implemented to mitigate such a trade-off but not used considering the computation overhead. In both datasets, *p*% is set to 6.25%, i.e. 1/16, which can be easily calculated with 4-bit right shifting as all numbers are in fixed-point representation.

*P* determines the adaptiveness of the system to the varying signal condition. The threshold can be less sensitive to signal varying if *P* is large, while it can be disturbed by the burst firing if *P* are too short. The frequent threshold updating also leads to more computation and higher power consumption. 3 s is used in both datasets.

The target detection count interval [*R*_2_, *R*_1_] is the most important parameter determining the acceptable detection rate range and eventually determines the saturated threshold level. There are several benefits to setting the target detection rate rather than conventionally deriving from multiple times to noise mean/median/root-mean-square/standard deviation values. 1) Compared to finding a multiplier, selecting one detection count interval can be more heuristic, building on previous neuroscience observations like [62] other than trial and error. 2) It guarantees the detection of useful information. It could happen using multiple times of signal statistics that the threshold can grow too high or low in some channels. That can lead to continuous detection or zero detection resulting in system failure. 3) The target detection rate range can potentially be used to improve the decoding performance. In more detail, if the resultant detection rate of a certain period approaches or exceeds *R*_1_, such period is more likely to be active, while if the resultant detection rate of another period is under *R*_2_, such period is more likely to be silent. The traditional method cannot provide such extra information (at least not in real-time), and we have proposed a Rate-Modulated LSTM based on such information, introduced in section 4.3.3, showing its superior performance. Notice that the target detection rate mentioned later will be the count interval upper limit *R*_1_ unless specified.

### 4.2. Spike count binning

The output of the spike detection is a binary stream. It has been found in [18, 45] that binning the binary threshold crossings into MUA counts at a certain period can improve the decoding accuracy. However, a trade-off between decoding accuracy and temporal resolution has to be made. In this work, a 50 ms bin period is used as in [18, 45].

### 4.3. Neural decoding

The binned MUA counts from different days are split into ten folds for decoding model cross-validation. For each validation cycle, one fold is used for training; one fold is used for validating the best model and the rest folds are used for testing. The average score across 10-fold validation is used as the final score of this day. Before feeding into the model, the binned MUA signals are normalised using the mean and standard deviation of the training set unless a special design is applied.

Two baseline decoding algorithms are implemented, namely Wiener Cascade Filter (WCF) and Long-Short Term Memory (LSTM), to decode the subject behaviour (finger velocity) from the spike events detected by the proposed spike detection algorithm.

#### 4.3.1. Wiener cascade filter

This adds non-linearity to the conventional Wiener filter [45, 63–65]. It consists of a linear Wiener filter and a static nonlinear system. The Wiener filter is trained by fitting the Lasso regression between the binned features of two recent time stamps and kinematic data with a regularisation factor of 0.01. A 3rd order polynomial between the Wiener filter output and the kinematic data is then fitted and cascaded after the Wiener filter to add non-linearity to the system.

#### 4.3.2. Long-short term memory

LSTM has been widely used in various machine learning problems, especially for time series analysis. Recently, it is also used in brain signal decoding achieving promising performance [6, 45, 66]. After model tuning, 150 LSTM cells are used with Adam optimiser at a learning rate of 0.0035 and batch size 64.

#### 4.3.3. Spike detection and neural decoding co-design

As mentioned before, the proposed spike detection algorithm can provide extraction information of the potential silent and active periods. We have designed a special input layer before the LSTM to utilise such extra information to modulate the binned spike count. A Rate-modulated LSTM (RM-LSTM) is proposed to have the capability to keep the consistent feature representation even though the input feature statistics are changed over time. The input layer consists of a firing rate modulation step and a dense neural network layer which is formulated as Equ.2

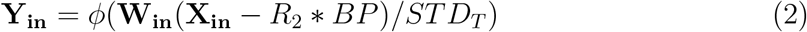

where **X**_**in**_ ∈ ℕ^*N*^ is the binned MUA signal, and N is the channel number. *STD*_*T*_ is the standard deviation of the training set, *R*_2_ is the target detection count lower limit, and BP is the bin period in seconds used for firing rate alignment. **W**_**in**_ ∈ ℝ^*N×N*^, *ϕ*(·) and **Y**_**in**_ ∈ ℝ^*N*^ are the weights, Sigmoid nonlinear activation and the output of the dense layer,

Consistent feature representation means the input of the LSTM can be at a similar scale statistically regardless of the varying of the binned MUA count statistics. This is achieved using the firing rate modulation. Input feature consistency is critical to maintaining the long-term stability of the decoding model, as the SNR can degrade with implant aging. Shifting the data via **X**_**in**_ − *R*_2_ ∗ *BP* makes the potential activate period positive while the silent period is negative. The input-output mapping of the model becomes zero-centred after shifting, as shown in Fig.1a firing rate modulation. Zero-centred mapping is easier to learn by the decoding model. At the same time, such mapping is input-feature-scale invariant. Even though the recording quality is degraded, the decoding model is supposed to retain its decoding performance. Moreover, such mapping is consistent for different target detection rates, and there is no need to retrain the decoding model after changing the threshold level of the spike detection algorithms.

The nonlinear response adds the capability of the models to simulate the nonlinear relationship between the input firing rate and the output that estimates behaviour. We assume the effect of the input firing rate on the behaviour (finger movement velocity) should show a saturation trend when the input firing rate is very high or small. Such a relationship can be well simulated using the Sigmoid function, a widely used activation function in neural networks.

Compared to the conventional LSTM, RM-LSTM used different input (**X**_**in**_ − *R*_2_ ∗ *BP*) and more parameters (**W**_**in**_). We built another two models: Rate-Subtracted LSTM (RS-LSTM), which only subtracts the rate from the input and Input-Dense LSTM (ID-LSTM), which only has the input dense layer and nonlinear activation. RS-LSTM controls the number of parameters to identify the improvement made from firing rate modulation, and ID-LSTM controls the input feature statistics to identify the contribution of the increased model size.

## 5. Results

### 5.1. Evaluation metrics

Several metrics have been used to evaluate the performance of the proposed spike detection and decoding algorithms.

For spike detection, Sensitivity (Sens), False Detection Rate (FDR) and Accuracy (Acc) are used and defined as below:

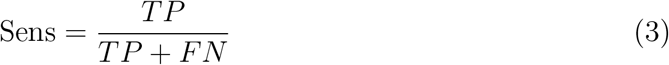

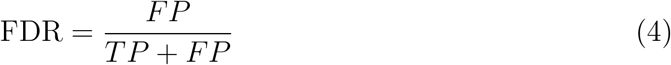

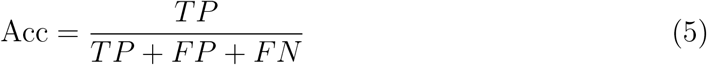

where TP, true positive, is the number of spikes that have been correctly detected; FP, false positive, is the number of periods that have been incorrectly detected as spikes; FN, false negative, is the number of true spikes that the algorithm has not detected.

Sens describes the ability of the algorithm to refine the spikes, while FDR evaluates the algorithm’s ability to reject noise. An algorithm cannot achieve the best performance on both metrics simultaneously, and Acc is the metric trading-off between them.

For neural decoding, the Parson Correlation Coefficient (CC) between the behaviour signal and predicted output is used and formulated as Equ.6.

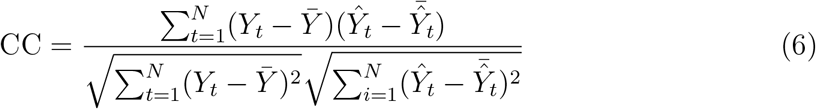

where *Y*_*t*_ and 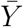 are the true target velocities at timesteps *t* and the average, *Ŷ*_*t*_ and 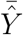 are the predicted velocities and the average. N is the total number of samples in this trial. CC is measured on both the x and y-axis and averaged across 10-fold validation.

### 5.2. Spike detection performance in synthetic dataset

We first tested the proposed spike detection algorithm on the synthetic dataset to quantitatively evaluate the algorithm compared to other state-of-the-art spike detection algorithms.

The performance at different noise levels are plotted in Fig.2a along with other algorithms. The proposed real-time low-complexity fixed-point multiplication-free algorithm has shown the highest detection accuracy at all noise levels among our previous work[35], offline spike detection[39], real-time float point algorithm[40] and fixed point algorithm[30]. It also performs better than the high complexity algorithm[67] in 0.05 to 0.15 noise levels. The proposed algorithm achieves nearly perfect detection at low noise levels while still above 90% accuracy at 0.2 noise level.

**Figure 2.**
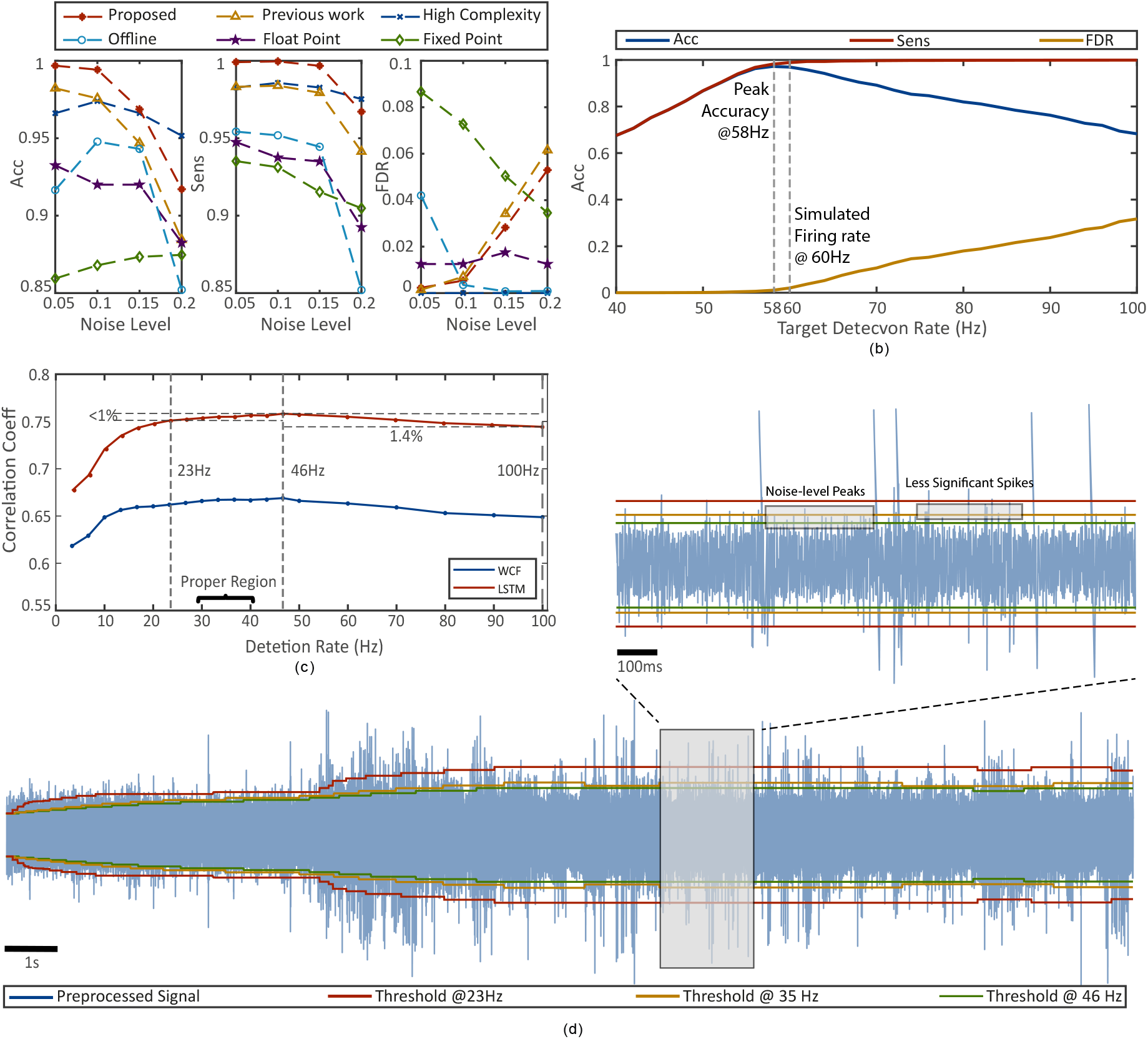
a) Spike detection accuracy, sensitivity, and false detection rate of the proposed algorithm compared to our previous work[35], and exemplar methods including a high complexity algorithm[67], an offline algorithm[39], a low complexity float-point algorithm[40] and a low complexity fixed-point algorithm[30]. b) The detection performance of the proposed algorithm when setting target detection upper bound at different values. The simulated recording consists of the spikes firing at 60Hz while the accuracy peaks at 58Hz. c) Decoding correlation coefficients using LSTM and WCF v.s. target detection rate. CC peaks at 46Hz while 30 - 40Hz should be the most reasonable rate to be detected. However, if double the detection rate to 100 Hz, 1.4% performance can happen while only less than 1% degradation can happen if the detection rate is halved 23Hz. d) Preprocessed signal with three different thresholds: 23 Hz, under-detected, only significant spikes are detected. 35 Hz, properly detection. 46 Hz, over-detected, many noise-level peaks are detected. Note that in practice, the absolute value is used instead of dual thresholds, but we kept the sign here for better visualisation.

To evaluate the performance of the proposed algorithm in practice quantitatively, we tried to decode both the threshold crossing derived from signal STD values and the result of the proposed algorithm. The average decoding CC of 24 days of recording is shown in Table. 1. The STD threshold is calibrated for each channel and each day. The proposed spike with a fixed threshold setting at a detection rate 46Hz achieves even higher decoding CC than the result of calibrated STD thresholding. There is only 1% CC improvement for fine-tuning the proposed algorithm (i.e. no need for fine-tuning). Compared to our previous work in [35] which has already achieved ultra-low resource usage and power consumption, we further reduce the resources used to less than one thrid of before, which directly translates to the area usage reduction of the ultimate ASIC design. A summary of the hardware cost and other specifications of the proposed algorithm is shown in Table. 2 along with other state-of-the-art algorithms.

**Table 1.** The decoding performance of WCF and LSTM using firing rate feature detected from RMS-based thresholding (provided by the public dataset and used by all other previous works) and the proposed algorithm. ”Calibrated” means the threshold settings are calibrated to achieve the best performance for each channel for every day.

|  | STD Calibrated | Proposed fixed at 46Hz | Proposed - Calibrated |
| --- | --- | --- | --- |
| WCF | 0.636 | 0.663 | 0.668 |
| LSTM | 0.738 | 0.753 | 0.759 |

**Table 2.** Hardware specification comparison of the proposed algorithm with other state-of-the-art implementations.

|  | Proposed | [35] | [68] | [24] | [69] | [40] |
| --- | --- | --- | --- | --- | --- | --- |
| Technology(nm) | 40<sup>1</sup> | 40<sup>1</sup> | 16<sup>2</sup> | 180 | 130 | 28 |
| Sampling rate(kHz) | 900 | 900 | 2560 | 800 | 800 | 10000 |
| Supply Voltage(V) | 1.8 | 1.8 | 3.3 | 1.8 | 1.2 | 0.8 |
| Channels | 128 | 128 | 128 | 128 | 64 | 1024 |
| Power per Ch.($\mu W$) | 0.21/0.03<sup>3</sup> | 0.29/0.04<sup>3</sup> | - | 0.64 | 3.04-4.54 | 1.45 |
| Logic Cells | 92 | 299 | 53607 | - | - | - |
| DSPs | 0 | 0 | 6 | - | - | - |
| Memory(KB) | 1.5 | 3 | 36 | - | - | - |
| Pre-emphasis | ADT | ADT | NEO | NEO | None | ADO-ASO |
| Thresholding | Firing rate | Firing rate | Mean | RMS | Mean | Median |

1 Implemented using reconfigurable logic (FPGA) platform Lattice ICE40LP1K
2 Implemented using reconfigurable logic (FPGA) platform Xilinx ZCU106
3 Incremental dynamic power

### 5.3. Target spike detection rate and spike detection performance

In order to find the relationship between the spike detection performance and neural decoding performance, we need a measure of the spike detection performance on the realistic dataset. It is challenging as there is no ground truth. However, with the proposed spike detection algorithm, we can perceive the spike detection performance using the target detection rate. More specifically, if we set the target spike detection rate close to the real, local firing rate, the algorithm should provide the highest spike detection performance. In order to prove that, we tried to use different target detection rates to set the threshold on the synthetic dataset composed of the spikes firing at 60Hz in total.

The detection accuracy, false detection rate, and sensitivity are shown in Fig.2b. The spike detection performance peaks at *R*_1_ = 56Hz, close to the 60Hz real spike firing rate. It demonstrates that the proposed spike detection algorithm can provide the highest detection performance when the target detection rate is close to the actual firing rate. It can also be observed that when the target detection rate is above the actual firing rate, both sensitivity and false detection rate increase, which means more spikes are detected while more noise peaks are recognised as spikes and vice versa.

### 5.4. The relationship between the spike detection and neural decoding performance

Utah array collected data that has been spike sorted [58] show 3 to 5 clusters per channel, whilst [70] suggests there are 3.8-4 clusters on average per channel when no spikes are discarded. Therefore, a reasonable assumption of the target detection rate should be 30-40 Hz for a ticker control task (as motor neurons can fire at below 10 Hz when operating little-effort tasks). The actual interval can be even lower as the number of neurons can be fewer than the observed clusters and the subject did not operate the ticker continuously. With this assumption, we are able to set a target detection rate interval to get the most accurate spike detection results on a realistic dataset for the motor tasks. Varying the target detection rate allows us to change the spike detection performance and builds the relationship between detection and decoding.

Fig.2c is obtained after decoding the binned spike detection results with WCF and LSTM. 30-40 Hz should be the desired detection rate achieving the highest detection performance, and is supposed to provide the best spike detection performance as well. However, the decoding performance of both methods peaks at 46Hz. Such unmatching suggests the nonlinear relationship between the detection and decoding.

This desired detection rate interval might be inaccurate. To be more rigorous, we also visually observed the spike detection result in Fig.2d. It shows a recording snapshot with the spike detection threshold generated with *R*_1_ = 23 Hz, 35 Hz and 46 Hz in red, yellow and green separately. The threshold at 46Hz is obviously adapted to the level only a little above the noise floor. Compared to the threshold at 35Hz, which is the best threshold to discriminate spikes from the noise by visual inspection, this lower threshold detected more noise-level peaks (boxed). Both theoretically speaking and by inspection, detection at 46Hz is not an ideal spike detection setting but achieves the highest decoding performance.

This finding does not match what is expected intuitively that better spike detection performance and better decoding performance. It suggests that some around-noise peaks are results in spikes actually firing far away. Detecting these spikes improves decoding performance despite also detecting more noise. This finding is also supported by the result in [70], suggesting that discarding the noise cluster is detrimental to the decoding performance in decoding neural signals using single-unit activities.

Another finding is that if the detection rate is doubled, to e.g. 100 Hz, which means about half of the spikes detected could be noise peaks, the performance is only degraded by about 1.4%. This suggests that the decoding models, even the simple filter-based model (WCF), are robust to noise. We only investigated a simple Long-Short-Term Memory neural network. It would be expected that if more advanced deep learning models are used, even better noise resistance can be achieved.

Finally, if the detection rate is halved to 23 Hz, many of the spikes with lower peak amplitudes (boxed). However, the decoding performance is only degraded by less than 1%. This finding is significant for on-implant spike detection, allowing us to send fewer data without sacrificing much performance. The threshold crossing can be represented using a binary stream at 1kbps. A lower detection rate means a more sparse signal, and the more sparse the signal is, the easier it is to be compressed. The data bandwidth can be even reduced using compression techniques[18], leading to less wireless transmission power.

Spike detection previously is an unsupervised problem in practice and highly depends on the results from synthetic data. Our finding is significant in providing a guide on setting the threshold. Especially for the brain machine interfaces with a limited power budget, the goal of spike detection would no longer be to find the best threshold level for the highest detection accuracy but the trade-off between the data bandwidth and decoding performance. With the firing-rate-based spike detection, the threshold level can be selected so it fulfils the system requirement on bandwidth and decoding accuracy without needing to consider the performance of the spike detection algorithm itself.

### 5.5. The performance of the co-design system

We used the target detection rate to bridge the neural decoding with spike detection. The improvement is shown in Fig.3a. Compared to the Conventional LSTM (Blue), over 1% improvement has been achieved using RM-LSTM (Red) at all different detection rates.

**Figure 3.**
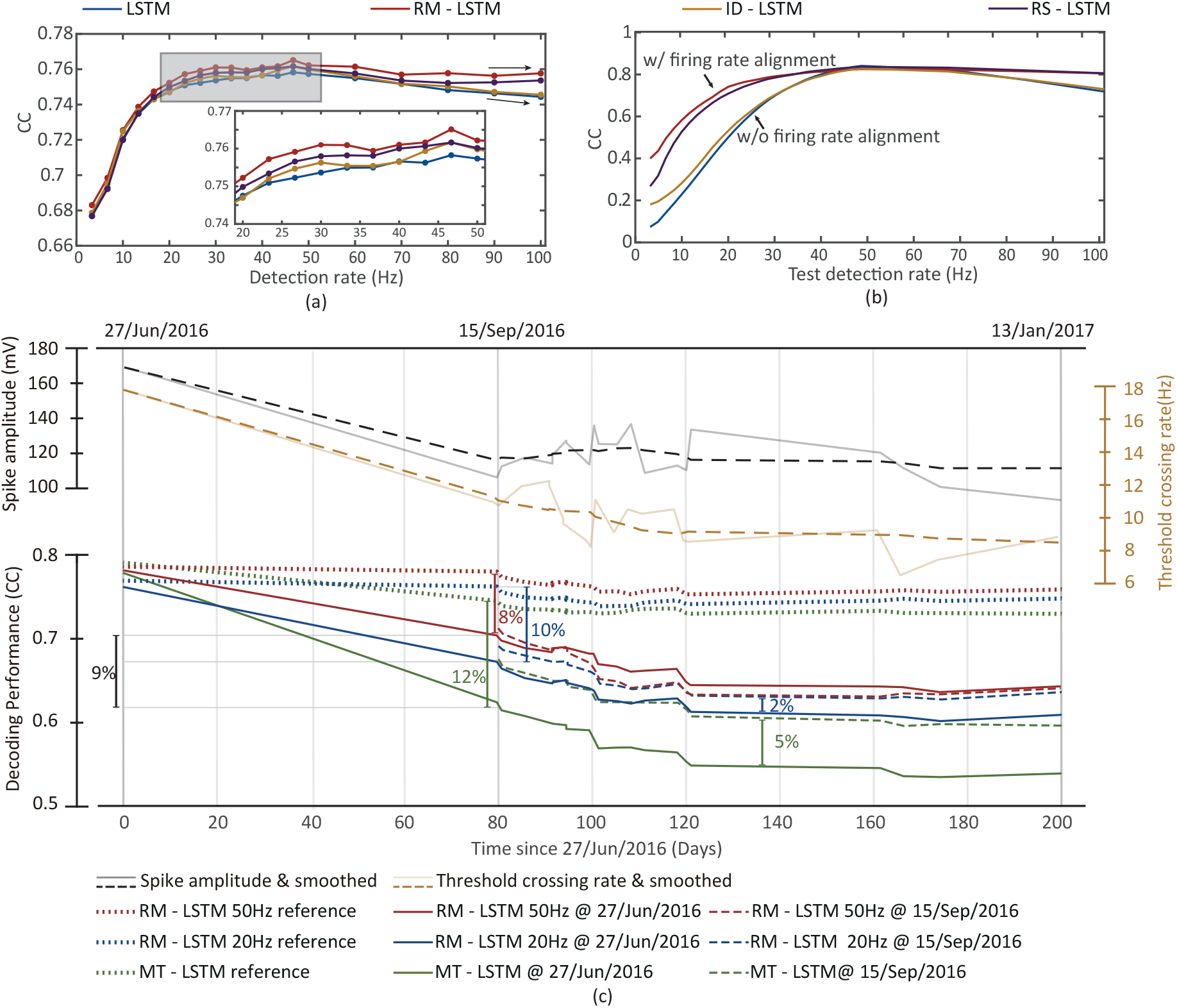
a) Decoding performance of different decoding models. RM-LSTM achieves the highest decoding performance among all models at all detection rates. b) The decoding performance when the model is trained with detections at 50 Hz and tested with detections at varying rates. The decoding performance is distinguishably improved when two rates are different by using the firing rate modulation. c) The spike amplitude and threshold crossing rate (STD threshold detected) over time, the long-time decoding performance over 200 days. RM-LSTM and MT-LSTM are trained on the data recorded on 27/Jun/2016 and 15/Sep/2016, simulating severe and mild SNR degradation with electrode aging after implementation. RM-LSTM is trained at a target detection rate of 20 Hz and 50 Hz, representing a compact setting with a threshold crossing rate close to MT-LSTM and a high-performance setting. Decoding performance curves are smoothed for better visualisation. The raw plot is given in the appendix

When the detection rate is above 40Hz, more noise will be detected. The detected noise will be added in MUA counts to the decoding model. Because of such noise, we can observe a decaying trend to the tail of the conventional LSTM. RM-LSTM however contains the decay suggesting the firing rate modulation (keeping consistent feature representation) not only helps to improve the model capability but also improves the model’s robustness to the noise.

To further understand such improvement in decoding performance, we designed the ID-LSTM model, which has the same number of parameters as RM-LSTM but without the firing rate modulation as Fig.1c shows. The yellow curve in Fig.3a shows its performance. ID-LSTM provides better decoding performance than conventional LSTM but is not as good as RM-LSTM. Such observation suggests that the improvement of RM-LSTM is not solely from the increased number of neural network parameters, but the firing rate information also matters.

We mentioned that firing rate modulation could improve the decoding performance because such an operation can keep a consistent feature representation and zero-centre the mapping, which is easier to learn. However, data normalisation can have a similar effect. In order to show that using the rate information of the proposed spike detection is better than the normalisation, we designed the RS-LSTM shown in Fig.1c shows. It subtracts the target rate instead of the mean calculated from the training set. It can be observed in Fig.3a that with such a simple subtraction, RS-LSTM achieves (purple) better performance than the ID-LSTM model, which has 48190 more parameters. We can also observe that the decaying trend with increased detection rate is also contained, similar to RM-LSTM.

Our results above not only demonstrates the superior performance of the proposed spike detection and neural decoding system co-design, but also provide an explanation as to where the improvement comes from, addressing the importance of consistent feature representation. It is essential that AI is not blinding applied but rather its application is informed by some knowledge of the “system”. We believe such a co-design system can thus provide better intuition in designing and optimising decodes performance than by simply applying complex deep learning architectures.

### 5.6. Long-term neural decoding performance

Maintaining long-term neural decoding performance is a challenging task in brain machine interfaces, because of the recording quality degrading with implant aging. At the top of Fig.3c, the black/grey curves show such decaying of the spike amplitude over 200 days. As the signal weakens, using traditional statistic-based spike detection results in fewer spikes being detected over time. As the yellow curves show, the average threshold crossing rate can drop from 18 Hz to below 10 Hz. (Spikes are detected using a threshold of 3.5 to 4 times the local STD values[61].)

Neural decoding is highly data-dependent. Maintaining consistent input features is one key to preventing the decoding performance from degrading. The conventional spike detection algorithm utilising a multiplier levelling up the noise statistics cannot provide such consistent features. Setting a smaller multiplier to lower the threshold level can resolve this issue to some extent, but manually tuning all channels to maintain a stable output stream is impractical.

The proposed spike algorithm can resolve this issue as the target detection rate guarantees a consistent threshold crossing rate to the decoding model over time, regardless of the noise levels and spike amplitude. Though more noise will be detected over time as the SNR degrades, this system co-design method is more robust to noisy input, as shown previously.

To evaluate the superior performance of the proposed system in long-term stability, we have tested three different models. In addition to the RM-LSTM and conventional LSTM, we implemented another model, Mean-Tracked LSTM (MT-LSTM). Its only difference from the conventional LSTM is that test data is normalised by the test data statistics rather than the values from the training set. This model simulates tracking of the input mean values, so-called Mean-tracked. The conventional LSTM and MT-LSTM use the threshold crossing of the STD threshold, while RI-LSTM uses the results of the proposed algorithm detection at 20 Hz and 50 Hz. The 20 Hz model keeps the threshold crossing rate close to the STD-detected ones, while the 50 Hz model allows further cell activities to be detected to improve decoding performance.

The decoding performance of different models is shown at the bottom of Fig.3c. We can clearly observe their decaying trends following the threshold crossing rate. The conventional LSTM cannot provide any long-term decoding ability because of the inconsistent threshold crossing rate over time. CC is consistently below 0.1 and therefore not plotted. RM-LSTM-50Hz (Red), RM-LSTM-20Hz (Blue) and MT-LSTM (Green) are trained with the data from different dates. The model trained on recordings collected on 15/Sep/2016 simulates the scenario when the electrode aging is mild (dash curves), while the one trained on 27/Jun/2016 simulates the scenario of severe electrode aging(solid curves). We also provide the results of the models trained and tested on the same day data as a reference observing how much the performance is degraded(dotted curves).

When the electrode ageing is mild, the nearly overlapped red and blue dash curves indicate that there is no significant difference using activities further away. Two RM-LSTM outperforms the MT-LSTM by about 2%. Such improvement is mainly contributed by the model capability instead of firing-rate modulation’s contribution to long-term stability because the RM-LSTM reference already outperforms MT-LSTM reference for about 2%. However, the superior performance of RM-LSTM is observed when the aging is severe. After 80 to 100 days of implementation, compared to the reference model, 12% performance degradation can happen on MT-LSTM and 10% for RM-LSTM-20Hz. However, it is only 8% for RM-LSTM-50Hz.

Moreover, comparing the severe electrode aging case to the mild aging one, there is about a 5% difference for MT-LSTM. However, it is only a 2% difference for RM-LSTM-20Hz and even better performance using RM-LSTM-50Hz (Because the data quality of 27/Jun/2016 is better than that of 15/Sep/2016). Compared to the performance of the first day of implementation, after 100 days of implementation, MT-LSTM degrades for more than 20% while both RM-LSTMs degrade for only around 10%. The co-design improves the the long-term decoding performance for nearly 10%.

Based on the observations above, it appears that consistent features and activities from further cells are both beneficial to improve long-term stability, especially when the electrode degradation is severe. The proposed RM-LSTM can take advantage of both, delivering superior long-term stability.

We also tested how the performance of RM-LSTM can be affected when fewer spikes are detected over time. We tried to train the model with the detection results in one target detection rate (50Hz) and tested at a different detection rate simulating more or fewer spikes will be detected as time goes by. We can observe the significant improvement in Fig.3b, especially when fewer spikes are detected when the rate information is used, demonstrating the robustness of the system co-design.

## 6. Discussions

Our results suggest that the relationship between spike detection and neural decoding performance is non-linear. The spike peaks of high amplitude contain substantial information to achieve acceptable decoding performance while detecting the spike peaks of amplitude comparable to the noise can improve the decoding performance despite more noise being detected. Such a finding brings opportunities for both brain machine interface engineering and neuroscience. Considering the proposed spike detection and neural decoding system co-design, we identify that maintaining the input data consistency is one key to improving the decoding performance and robustness, especially in the long term, when recording quality is degraded gradually with electrode ageing.

### 6.1. Opportunity for designing low power brain machine interfaces

In [16], It is suggested that relaxing some of the hardware requirements, for example, sampling frequency, amplifier performance, and ADC precision, can reduce the system power consumption by an order of magnitude without much degrading the decoding performance. It also shows that randomly missed spikes can only degrade decoding performance by a small amount. Until this work, there has not been reported any spike detection algorithm that can “miss” spikes heuristically.

In this work, we suggest relaxing the amount of data transmitted. Only transmitting the high-level peaks can reduce half of the data to be transmitted with less than 1% decoding performance degradation for behaviour decoding task. Half bandwidth reduction is significant in reducing transmission power, especially when wireless transmission is involved.

The threshold crossings can be encoded into a binary stream or the number of spikes detected in a certain period. The reduced detected spikes rate can either create a more sparse binary stream or reduce the dynamic range of the possible spike count range.

In both cases, the bandwidth can be reduced. Taking the latter case as an example, we assume the detection stream is binned at *τ* sec, and the detection rate is *λ* Hz following Poisson distribution. The number of arrival *N* of one bin is a Poisson process as Equ.7 shows.

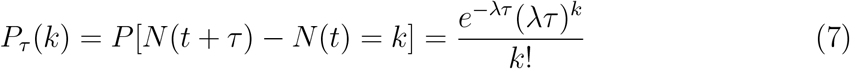

In practice, the spike count is saturated at a value *S* to limit the spike count data width. The probability of the spike count exceeding *S* (inclusive) is given in Equ.9 and it needs *L*^*S*^ bits for representing *S* values

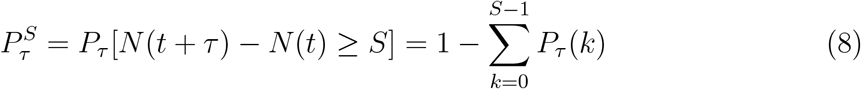

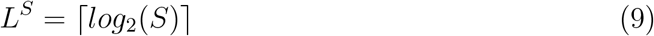

If we saturate the spike count at *P*_*τ*_ (*S*) *<* 1‰, when a 50Hz detection stream is binned at 0.01s, S is 6 with 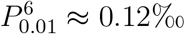. When the detection rate is reduced to 25Hz, S becomes 4 with 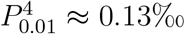. Therefore 3 bits are required for 50Hz spike detection counts, while it is 2 bits when detection at 25Hz.

From an entropy perspective, the theoretical minimum entropy coding length *L*_*τ*_ that losslessly compresses such a stream can be calculated as Equ.10.

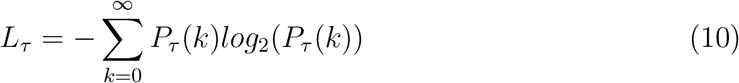

Fig.4 shows how the number of bits and minimum code length for representing the spike counts. The minimum coding length is 1.34 at 50Hz, while it becomes only 0.89 at 25Hz. 33% bandwidth can be reduced while it only introduces less than 1% decoding degradation, as Section.5.4 shows.

**Figure 4.**
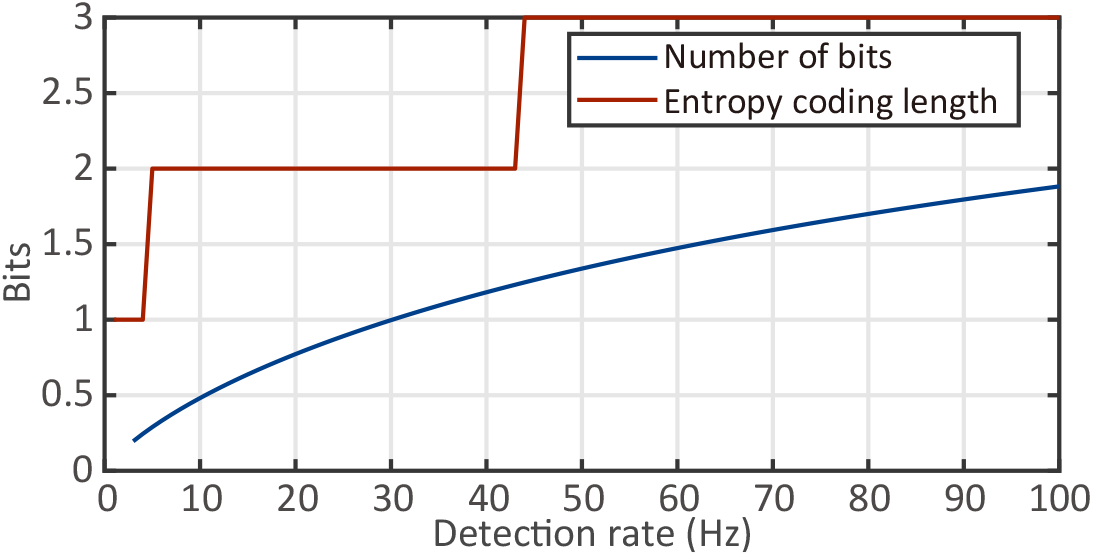
The number of bits and entropy encoding length for representing the spike counts

### 6.2. Feature consistency

Inconsistent features can degrade the performance of the learning models. This inconsistency can come in three different forms. The first form is invalid features. For example, the model is fed with Motor Cortex recordings for behaviour decoding but tested with Visual Cortex recordings, which is technically incorrect. The second form is the incomplete feature set. For example, the model is trained on neural recordings for reaching tasks but tested to decode placing tasks. Such inconsistency can be overcome by designing more generalised models.

More importantly, the third form is the statistical difference, which we can meet in long-term neural decoding. The recorded amplitude of action potentials can gradually decrease over 37% within two months after implantation[12]. Some studies [12, 45, 70] suggest that threshold crossing can achieve better long-term stability than using spike waveforms or spike sorting results, but do not provide an underlying reason.

We need to know how machine learning or deep learning models are trained to understand such a reason. There is a normalisation step before feeding input features into models and guaranteeing the input is not statistically changed. However, tested datasets can only be normalised using training dataset statistics as the tested dataset should be unknown to the model. Normalising the test data using training statistics can be a problem in long-term decoding. The realistic mean of the input models can be lower than the features used in training. Normalisation can shift the testing features’ centre to negative while it is expected to be at zero. That is why long-term degradation can happen (in addition to more noise being contaminated).

Using spike waveform as the feature can be affected the most as it is most sensitive to such a normalisation error. Using SUA from spike sorting has worse long-term stability than using MUA. Besides fewer spikes appearing in each cluster, the inconsistent spike waveform can negatively impact the clustering algorithm. Such artefacts will propagate to the decoding model and even lower the decoding accuracy. Spike detection is only affected by the reduced number of detections.

Knowing the reason for long-term stability degradation, one key to resolving this issue is maintaining the consistent input feature of the decoding model. The proposed spike detection and neural decoding co-design system addresses that. This idea can potentially be explored in other domains, such as spike waveforms, single-unit activity, or local field potentials.

### 6.3. Limitation of this work

The results of this work have been validated on 5 hours of recording over 7 months. However, the recordings are from a single subject operating one type of task. Findings are currently only valid for motor cortex behaviour decoding. The cross-subject generalisation is another important area to be investigated. Unfortunately, we cannot test so limited by the available data. We can expect improved cross-subject decoding performance if our claim on feature consistency is established.

## 7. Conclusion

In this work, we presented a methodology for BMI signal processing system co-design that significantly improves long-term decoding performance compared to current methods. By modulating the input feature with the target detection rate, we were able to improve decoding performance by more than 10% after 80 days of operation. We also demonstrated the nonlinear relationship between spike detection and decoding performance. Using the proposed co-design methodology, by reducing spike detection sensitivity to detect approximately half the spikes, results in only a 1% decoding degradation whilst also reducing communication bandwidth by at least 30%.

For the first time, this work tightly links spike detection with decoding. We believe the significant improvement presented in this work can motivate hypothesis-led data science strategies to further develop brain machine interfaces.

## Acknowledgement

The authors would like to thank Rodrigo Quian Quiroga (Leicester) and Nick Steinmetz (University College London) for making their datasets publically available.

